# Selection dynamics in transient compartmentalization

**DOI:** 10.1101/258269

**Authors:** Alex Blokhuis, David Lacoste, Philippe Nghe, Luca Peliti

## Abstract

Transient compartments have been recently shown to be able to maintain functional replicators in the context of prebiotic studies. Motivated by this experiment, we show that a broad class of selection dynamics is able to achieve this goal. We identify two key parameters, the relative amplification of non-active replicators (parasites) and the size of compartments. Since the basic ingredients of our model are the competition between a host and its parasite, and the diversity generated by small size compartments, our results are relevant to various phage-bacteria or virus-host ecology problems.

PACS numbers: 05.40.-a, 87.14.G-, 87.23.Kg

---

A central issue in origin of life studies is to explain how replicating functional molecules could have appeared and evolved towards higher complexity [1]. In 1965, Spiegelman showed experimentally that RNA could be replicated by an enzyme called *Qβ* RNA replicase, in the presence of free nucleotides and salt. Interestingly, he noticed that as the process is repeated, shorter and shorter RNA polymers appear, which he called parasites. Typically, these parasites are nonfunctional molecules which replicate faster than the RNA polymers introduced at the beginning of the experiment and which for this reason tend to dominate. Eventually, a polymer of only 218 bases remained out of the original chain of 4500 bases, which became known as Spiegelman’s monster. In 1971, Eigen conceptualized this observation by showing that for a given accuracy of replication and relative fitness of parasites, there is a maximal genome length that can be maintained without errors [2]. This result led to the following paradox: to be a functional replicator, a molecule must be long enough. However, if it is long, it can not be maintained since it will quickly be overtaken by parasites. Many works attempted to address the puzzle as reviewed in Ref. [3]. In some recent studies, spatial clustering was found to promote the survival of cooperating replicators [4–6]. This kind of observation is compatible with early theoretical views [7, 8], that compartmentalization could allow parasites to be controlled.

Small compartments are ideal for prebiotic scenarios, because they function as micro-reactors where chemical reactions are facilitated. Oparin imagined liquid-like compartments called coacervates, which could play a central role in the origin of life [9]. Although experimental verification of the prebiotic relevance of coacervates or other sorts of protocells remained scarce for many years, the idea has resurfaced recently in various systems of biological interest [10, 11]. An important aspect of the original Oparin scenario which has not been addressed in these studies is the possibility of a transient nature of the compartmentalization. In the present paper, we revisit group selection with transient compartmentalization. We were motivated by the relevance of transient compartmentalization in several scenarios [12–16] for the origin of life and by a recent experiment, in which small droplets containing RNA in a microfluidic device [17] were used as compartments. In this experiment, cycles of transient compartmentalization prevent the takeover by parasitic mutants. Cycles consist of the following steps: (i) inoculation, in which droplets are inoculated with a mixture of RNA molecules containing active ribozymes and inactive parasites, (ii) maturation, in which RNA is replicated by *Qβ* replicase, (iii) selection, in which compartments with a preferred value of the catalytic activity are selected, (iv) pooling, in which the content of the selected compartments is pooled. This protocol does not correspond to that of the Stochastic Corrector model [7] because of step (iv), which removes the separation between individual compartments.

The absence of parasite takeover was successfully explained in ref. [17] by a theoretical model which described the appearance of parasites within a given lineage as a result of mutations during the replication process. In this work we wish to account for these observations in a more general sense. We show that the value of the mutation rate does not play an essential role as long as it is small [18], and that the entire shape of the selection function is not needed to describe the fate of the system.

Let us consider an infinite population of compartments. Each compartment is initially seeded with *n* replicating molecules, where *n* is a random variable, Poisson distributed with average equal to λ. In addition, each compartment also contains a large and constant numbers of enzymes, *n*_*Qβ*_ and of activated nucleotides *n_u_.* Among the *n* replicating molecules, *m* are ribozymes, and the remaining *n — m* are parasites. Let *x* be the initial fraction of ribozymes and 1 — *x* that of parasites. After this inoculation phase, compartments evolve by letting the total number of molecules grow by consuming activated nucleotides.

In practice, the time of incubation of the compartments is fixed and longer than the time after which activated nucleotides become exhausted. The kinetics is initially exponential because the synthesis of RNA is autocatalytic at low concentration of templates. Therefore, the average number 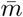 of ribozymes and 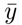 of parasites grow according to

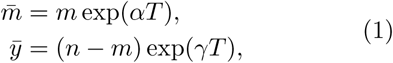

where *T* denotes the time and *α* (resp. *γ*) denote the average growth rate of the ribozymes (resp. parasites) during this exponential growth phase. The relevant quantity for this dynamics is the ratio of the number of daughters of one parasite molecule and that of the daughters of one ribozyme molecule: Λ = exp((*γ — α)T*)). Note that Λ > 1 since *γ > α*. This exponential growth phase (maturation phase) ends, when the total number of templates 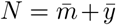 reaches the constant value *n*_*Qβ*_ which is the same for all compartments. After this point, the kinetics switches to a linear one, because enzymes rather than templates are limiting [19]. Importantly, during this linear regime the ratio of ribozymes and parasites

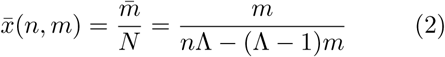

does not change. Apart from neglecting very small fluctuations in and *n_u_*, our assumption that *N* is constant means that the effects of fluctuations of growth rates of both species and the effect of a possible dependence of Λ on *m* and *n* are not considered. These two stochastic effects have been modeled in detail in the Supplemental Materials [18]. In the end, we find that they do not alter significantly the predictions of the present deterministic model for the conditions of the experiment.

Two types of parasites can appear: *hard parasites* are formed when the replicase overlooks or skips a large part of the sequence of the ribozyme during replication. The resulting polymers are significantly shorter than that of the ribozyme and will therefore replicate much faster. Based on the experiments of [17], we estimated Λ to be in a range from 10 to about 470, as explained in the Supplemental Materials [18]. In contrast, if the replicase makes errors but keeps overall the length of the polymers unchanged, then the replication time is essentially unaffected. In that case, one speaks of *soft parasites,* and the corresponding Λ is close to unity. It is important to appreciate that the distinction between *hard* or *soft parasites* is not only a matter of replication rates, because Λ also contains the time *T*, so depending on both parameters, parasites could be classified as either *hard* or *soft.*

The compartments are then selected according to a selection function f (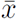) ≥ 0. A specific form which is compatible with [17] is the sigmoid function

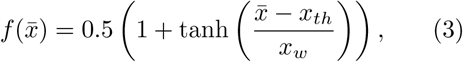

with *x_th_* = 0.25 and *x_w_* = 0.1. Note that this function takes a small but non-zero value for 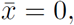, namely 0.5(1 —tanh(*x_th_/x_w_*)) = 0.0067, which represents the fitness of a pure parasite compartment. This is in contrast with the linear selection function chosen in a recent study of a similar system [20].

After the selection phase, the resulting products are pooled and the process is restarted with newly formed compartments. We wish to evaluate the steady-state ratio *x* of ribozymes when many rounds of the process have taken place. The probability distribution of the initial condition (*n, m*) is given by

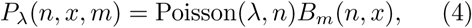

where *B_m_(n, x)* is the Binomial distribution for *m* ∊ {0,…, *n*} of parameter *x* ∊ [0,1]. The average of 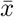 after the selection step is given by

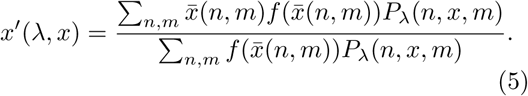

The sty-state value eadof *x* is the stable solution of

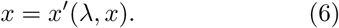

It is easier to evaluate *Δx = x′(λ, x) − x* as a function of λ. The steady-state value corresponds to the line *Δx* = 0 separating negative values above from positive values below as shown in Fig. 8.

We construct a phase diagram in the (λ, Δ) plane, by numerically evaluating the bounds of stability of the fixed point *x* = 0 from the condition:

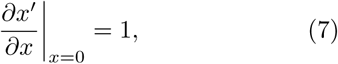

and similarly for the other fixed point *x* = 1. The resulting phase diagram, as shown in Fig. 10, shows four distinct phases. In the orange (resp. light blue P region) region R, the only stable fixed point is *x* = 1 (resp. *x* = 0). In the green region, *x* = 0 and *x* = 1 are both stable fixed points. The system converges towards one fixed point or the other depending on the initial condition: for this reason we call this region B for bistable. In the violet region, *x* = 0 and *x* = 1 are both unstable fixed points, but there exists a third stable fixed point *x\** with 0 < *x\** < 1. We call this a coexistence region (C). All of these phases can be seen in Fig. 8. In the Supplemental Materials [18], we discuss other aspects of the phase behavior which are not captured by this treatment. We also show there that many features of this phase diagram remain if a linear selection function is used instead of Eq. (3)

It is interesting to analyze separately some specific limits for which the asymptotes of the phase diagram can be computed exactly. Let us consider

- λ >> 1: bulk behavior
- Λ >> 1: *hard* parasites
- Λ close to 1: *soft* parasites

For large λ, we can neglect the fluctuations of *n,* i.e. the total number of replicating molecules (ribozymes plus parasites) in the seeded compartment. Indeed, *n* is Poisson distributed with parameter λ, therefore *Var(n)/λ^2^* = 1/λ << 1. For large λ, Λ close to 1 and *x* close to 1 (resp. 0), the most abundant compartments verify *m = n* or *m = n* — 1 (resp. m = 0 or m = 1). By considering only these compartments in the recursion relation [18], one finds that the condition of stability of the fixed point *x* = 0 leads to

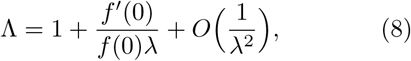

for an arbitrary selection function and Λ ≃ 1 + 19.86/λ for the selection function of Eq. (3). This equation indeed characterizes the separation between the parasite and coexistence regime at large λ in Fig. 10. A similar equation is found for the fixed point at *x* = 1

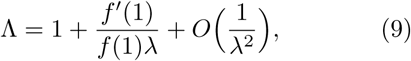

yielding Λ ≃ 1+6.12 10^−6^/λ for this selection function for the separation between ribozyme and coexistence regions. For Λ close enough to 1, we have a ribozyme phase. The asymptotes given by (8) and (9) border the coexistence region in Fig. 10. This supports the observation that *soft* parasites can coexist with ribozymes.

Let us now study the *hard* parasite limit, namely Λ ≫ 1, and finite λ. In this regime, we only need to consider three types of compartments: compartments made of pure ribozymes, such that *m = n ≠ 0*, compartments containing parasites, and empty compartments, i.e. such that *n* = 0. One can introduce three inoculation probabilities for these cases *Pribo, Ppara*, and *Pzero*. Using Eq. (4), one finds

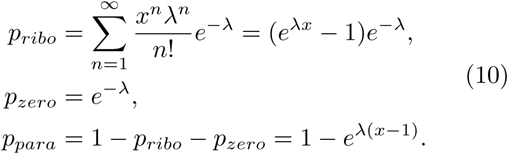

Assuming that in compartments containing parasites they will overwhelm the ribozymes, and inserting these values in (18), we find

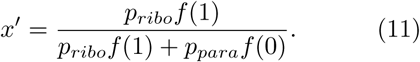

Evaluating the fixed-point stability of *x* =1 using (7), we find that the boundary value of λ satisfies

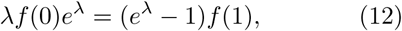

for an arbitrary selection function. A similar calculation at the ixed point *x* = 0 leads to the other vertical separation line given by

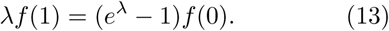

The solution of Eq. (12) (resp. Eq. (13)) is λ ≃ 149.41 (resp. λ ≃ 6.95) which compare well with the vertical separation lines in Fig. 10.

In ref. [17] a comparison was made of the system behavior as a function of the number of selection rounds in three possible protocols: (i) No compartments (bulk behavior), (ii) compartments with no selection, (iii) compartments with selection. Such a comparison based on our theoretical model is shown in Fig. 3 for parameter values corresponding to the coexistence region of Fig. 10. As expected, the fraction of ribozymes decreases towards zero rapidly in case (i), and somewhat less quickly in case (ii). Only in case (iii) is it possible to maintain a non-zero ribozyme fraction on long times. It is indeed observed that the ribozyme fraction eventually vanishes for protocols (i) and (ii) in the experiment of Ref. [17]. In case (iii), a decrease of the ribozyme fraction is observed. The last two points of figure 2C (top panel) in this reference are an indication that the system may eventually reach ribozyme-parasite coexistence in this regime.

**Fig. 1.**
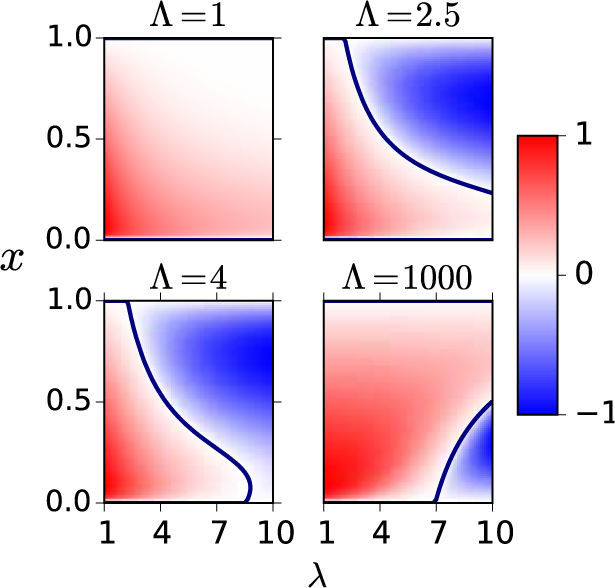
Contour plots of *Δx* for four values of A = 1,2.5,4 and 1000 in the plane (*x, λ*), with red (resp. blue) regions corresponding to *Δx* > 0 (resp. *Δx* < 0).

**Fig. 2.**
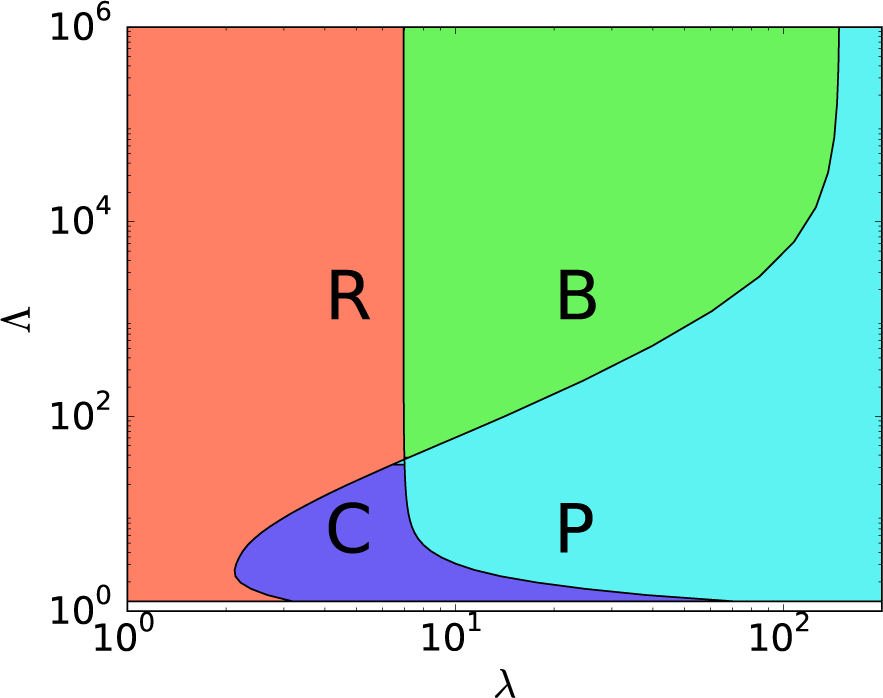
Phase diagram of the transient compartmentalization dynamics with the selection function of Eq. (3) in the (λ, Λ) plane. The phases are: *R*: pure Ribozyme, B: Bistable, C: Coexistence, P: pure Parasite.

**Fig. 3.**
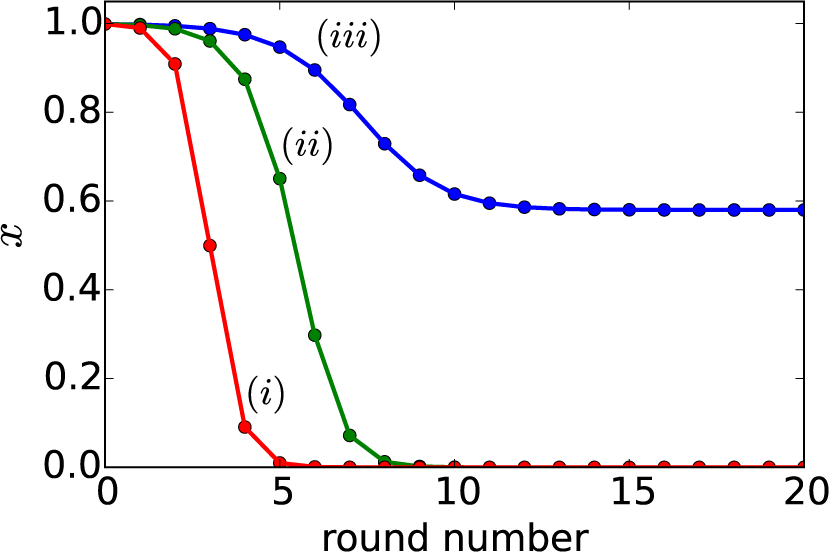
Evolution of the average ribozyme fraction *x* as function of the number of rounds for the three protocols, namely (i) No compartments (bulk behavior), (ii) compartments with no selection, (iii) compartments with selection. We choose λ = 5 and Λ = 10, corresponding to the coexistence region of Fig. 10.

In figure 4 we show the behavior of the distribution of the ribozyme fraction after the growth phase, i.e. 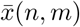(defined in Eq. (17)) as a function of round number. The parameters are Λ = 5 and λ = 10, corresponding to the parasite region, where the final state of the system is *x* = 0, and the initial condition is *x* = 0.999. Note that the distribution of 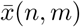 is discrete, since many values are not accessible in the allowed range of *n* and *m*. At *t* = 0, it exhibits a sharp peak near *X* = 1 coexisting with a broad peak at small values of *X*. As time proceeds, the weight of the distribution shifts to the peak at small values of *X*, since in this case selection is not sufficiently strong to favor the peak near *X* = 1 and parasites eventually take over.

**Fig. 4.**
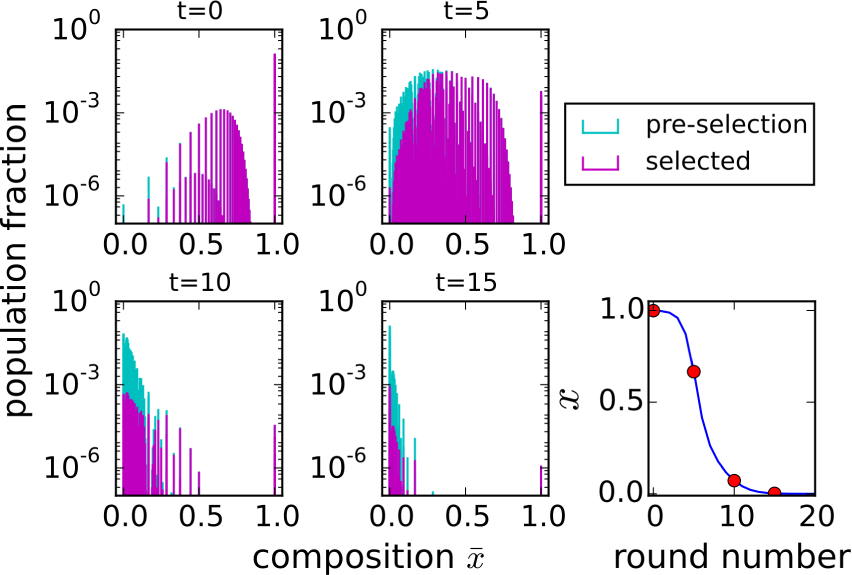
Evolution of the distributions of ribozyme fraction 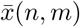 before and after selection at different times. The chosen times are shown as red circles in the lower right panel, which represents the evolution of the average fraction *x* as a function of the number of selection rounds.

In conclusion, we captured the behavior of transient compartmentalization with a model containing only two parameters, which remarkably suffices to capture the main features of the transient compartimentalization experiment [17]. The model predictions are summarized in a phase diagram, which has been derived for an arbitrary selection function.

Given its basic ingredients, the competition between a host and its parasite, and the diversity generated by small size compartments, which is required for selection to be efficient [21], the model has broad applicability. It could for instance be relevant for phage-bacteria ecology problems, which typically experience a similar life cycle of transient replication in cellular compartments during infection [22].

Our work also clarifies that group selection is able to purge the parasites even when compartments are transient. If such dynamics of compartments is applicable to protocells [9], the mechanism discussed here could represent an important element in scenarios on the origins of life.

A.B. was supported by the Agence Nationale de la Recherche (ANR-10-IDEX-0001-02, IRIS OCAV). L.P. acknowledges support from a chair of the Labex CelTisPhysBio (ANR-10-LBX-0038). He would like to thank ESPCI and its director, J.-F. Joanny, for a most pleasant hospitality.

## Supplementary Material

Here we provide more details on (i) the determination of the parameter Λ from experimental data and its intercompartment variation, (ii) the role of growth rate fluctations, (iii) the impact of mutations in the limit Λ → ∞, (iv) details on the derivation of the asymptotes, for λ → ∞ and Λ ≃ 1, (v) further aspects of the phase diagram, and (vi) a comparison of the phase diagram for linear and non-linear selection functions.

### I. Exponential Growth and the Value of Λ

The following table contains experimental values measured in Ref. [17] for the ribozyme and three different parasites. The nucleotide length, doubling time (*T_d_*), and relative replication rate (*r*), are reported, from which we infer Λ in the final column. The doubling time *T_d_* for the ribozyme is related to the growth rate *α* introduced in the main text by *T_d_* = ln(2)/α, and similarly the doubling times of the parasites is related to the γ by *T_d_* = ln(2)/γ.

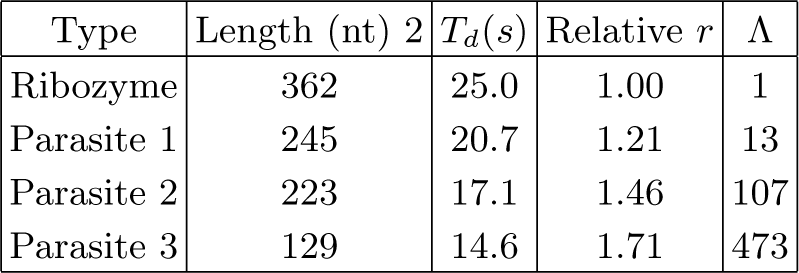

In the experiment, a typical compartment contains λ RNA molecules that can be ribozyme or parasite, 2.6-10^6^ molecules of Qβ replicase, and 1.0 · 10^10^ molecules of each NTP. Replication takes place by complexation of RNA with Qβ replicase, which uses NTPs to make a complementary copy. This copy is then itself replicated to reproduce the original. There is a large amount of nucleotides, so that exponential growth of the target RNA proceeds until N ≈ *n*_*Qβ*_. Starting from a single molecule, it takes *nD* = log_2_ *n* _*Qβ*_ = 21.4 doubling times to reach this regime. In a parasite-ribozyme mixture, we can estimate Λ using the relative *r*:

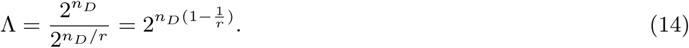

Another important assumption of our model, is that we neglect a possible dependence of Λ on *n* and *m*. In order to test this assumption we have estimated the fluctuations of Λ in the following way. We recall that the total number of RNA at the end of the exponential phase is the constant *n*_*Qβ*_ given above, thus *N* = (*n* — *m*)2*^nD^* + *m2^nD/r^* = *n*_*Qβ*_. We first solve for *n_D_* in this equation for a given *n* and *m* and then we use this result into Eq. (14) to obtain Λ for a given *n* and *m*. We show in figure 5 a typical plot of the values taken by this function Λ(*n, m*) for a particular choice of n and m, together with the probability distribution *P\λ(n,x,m)* defined in Eq. (4) of the main text. In general Λ(*n,m*) is close to a constant for soft parasites (*r* = 1.2), and is less constant for hard parasites (*r* = 1.7). Even in the later case however, A hardly varies in the range of *n,m* values where the probability distribution takes significant weight. We conclude that the assumption of neglecting a possible dependence of Λ on *n* and *m* has only a minor effect on our results.

**Fig. 5.**
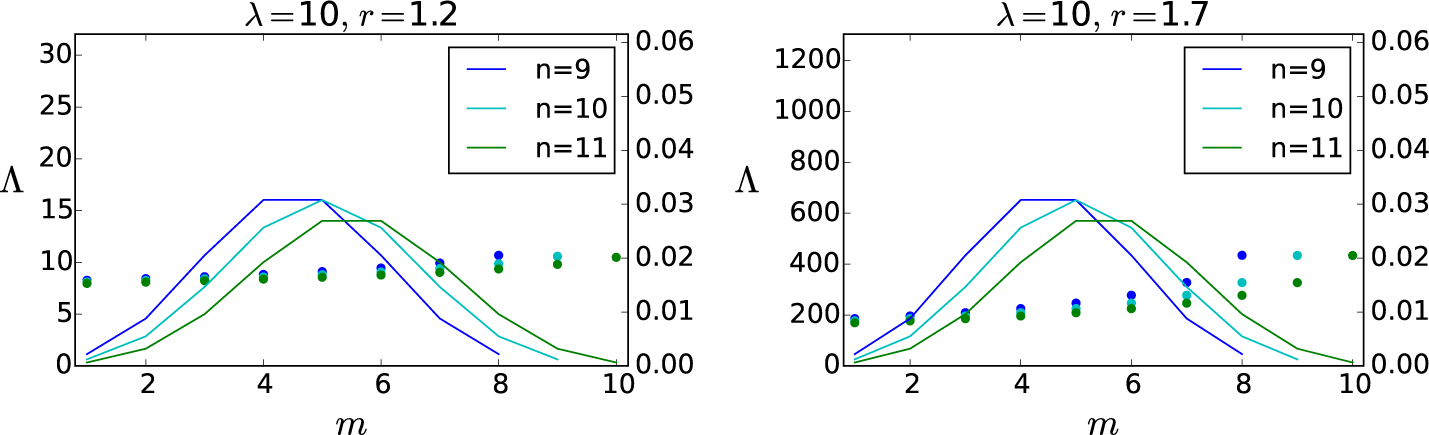
Plots for Λ (colored dots) as function of the parameters (*n, m*) characterizing the initial composition of the compartments relative growth rates *r* and *λ* = 10, together with the probability distribution *P_λ_* (*n, x, m*) (solid lines) for *x* = 0.5.

### II. Growth Rates Fluctuations

The model presented in the main text is purely deterministic, but fluctuations are still present due to the initial condition. Since growth rates typically depend on the initial condition, they will fluctuate when sampling the initial condition. While these fluctuations are already present in a deterministic model, effects associated to other fluctuations in the growth rates, which may occur at time scales smaller than the duration of the growth phase require a stochastic approach. Replication is intrinsically stochastic and therefore this additional source of fluctuations in grow rates could be present. In order to estimate such an effect, we have implemented below a stochastic version of our model.

A stochastic component in the growth phase of the ribozymes and parasites can be included using a discrete Langevin approach. In such an approach, Eq. (1) is modified to become:

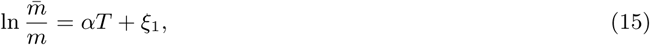

where the first term on the right hand side accounts for the deterministic contribution we had before and *ξ*_1_ is a Gaussian random variable of mean zero and variance *σ_1_.* Similarly for the parasites:

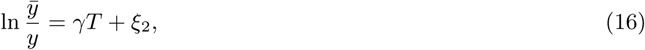

where ξ_2_ is another similar noise controlling the growth of the parasites.

The noise which has been introduced here could describe either fluctuations of growth rate *α* or of the duration of replication. Note also that we still define Λ with respect to the mean time and growth rates as Λ = exp((γ − α)T)). Since *N* is fundamentally fixed by the number of enzymes *n*_*Qβ*_ in this problem, we still assume that *N* is fixed in this stochastic model. Then this condition N = 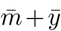 leads to a constraint between the noise ξ_1_ and ξ_2_, which means that these two noises must be correlated. Then, Eq (2) is modified as

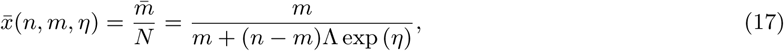

where we have introduced the random variable n = ξ_2_ — ξ_1_. Let us introduce the variance of η which we call σ^2^. This is the main parameter controlling the growth noise. Eq. (5) is now modified as follows

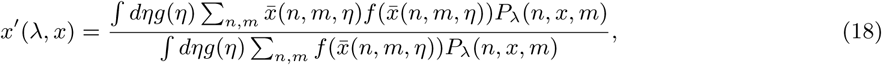

where g(η) is the Gaussian distribution of mean zero and variance σ^2^. In order to evaluate the correction due to the noise η, we expand the integrand in the numerator and denominator with respect to n and we perform the Gaussian integrals. The result is a modified recursion relation which contains a correction term proportional to σ^2^. The explicit expression of this correction is lengthy and not given here, it was evaluated numerically.

This Taylor expansion is justified if the amplitude of the noise σ^2^ is sufficiently small. In order to assess this, we investigate the various sources of noises in this problem. The noise could be due to the arrival times of *Qβ* or from the replication process itself. For the first source of noise, the time scale to form an *Qβ* − RNA complex due to diffusion can be evaluated as 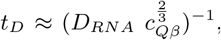, where D_RNA_ is the diffusion constant for an RNA strand (length ≈ 300) and *c*_*Qβ*_ the concentration of *Qβ* replicase. We found this timescale to be over 2 10^4^ times smaller than replication times (15-25s, see SM 1). Due to this large difference in timescales, the noise in this problem should primarily be caused by the replication rather than by the binding of a Qβ to an RNA strand. Let us now look at the noise due to replication. Once a Qβ enzyme is bound to a single RNA molecule, the total replication time *τ* can be written as a sum of the dwell times of all the nucleotides to be added to the template and which are themselves exponentially distributed. When the binding rates are identical, the resulting distribution of *τ* is a gamma distribution with a coefficient of variation 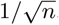, in terms of *n* the number of nucleotides as shown in D. Floyd et al. (2010). Since our replicating molecules are long, typically between *n* ≃ 150 to 300, this coefficient of variation is quite small. This coefficient of variation is expected to increase due to the number of doubling times in the replicating phase, which is typically of the order of 20. In the end although we can not provide an accurate estimate of the noise of *η*, all these factors suggest a small noise amplitude for σ^2^.

In the worst possible case, we would have σ^2^ = 1 which is the case shown in the two figures below. The red contour plots of *Δx* corresponding to that of Fig 1 of the manuscript, the blue ones correspond to the prediction of a stochastic version of the same model including the correction due to σ^2^. We only show the plots for Λ = 4 (left) and Λ = 2.5 (right), because we find that for high values of Λ the noise has only a negligible effect even in this worst case scenario, which is reasonable. While we see that the noise *η* affects significantly the contour plots in this worst case scenario, when σ^2^ = 1, the effect is quite small with a more realistic estimate of the noise namely σ^2^ =0.1 as shown in figure 7:

All these results support our view that while the growth is intrinsically stochastic the deterministic model we have developed with *a* constant *N* and fluctuations only in the initial conditions indeed capture the main features of the experiment we are interested in. As shown in figures 6 and 7, the stability of the ribozyme phase in the new stochastic model is enhanced with respect to the deterministic model. This confirms that fluctuations are essential to stabilize the ribozyme phase, and favor it whether they come from the initial condition as in the deterministic model or from other sources as in the stochastic developed here.

**Fig. 6.**
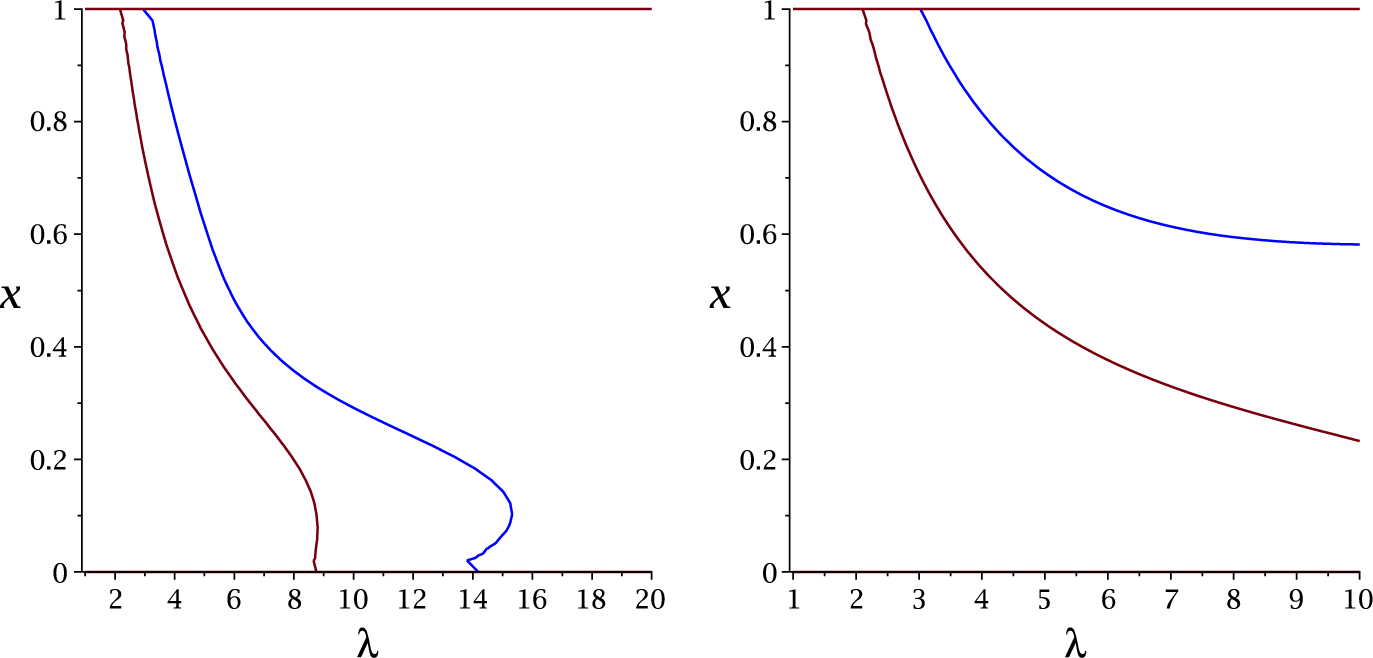
Contour plots of *Δx* similar to that of Fig 1 of the manuscript, with no correction due to noise (red solid line) and including corrections due to noise with *σ*^2^ = 1 (blue solid line). The figures correspond to Λ = 4 (left) and Λ = 2.5 (right).

**Fig. 7.**
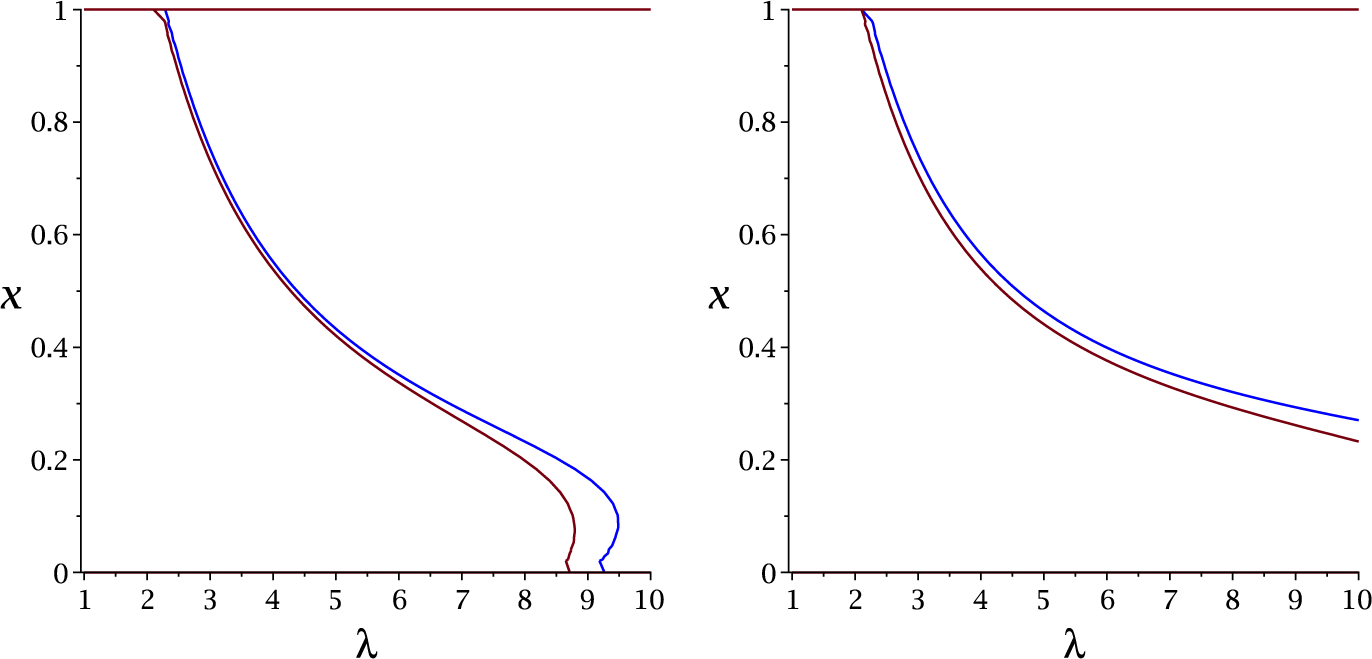
Contour plots of *Δx* similar to that of Fig 1 of the manuscript, with no correction due to noise (red solid line) and including corrections due to noise with *σ^2^* = 0.1 (blue solid line). The figures correspond to Λ = 4 (left) and Λ = 2.5 (right).

### III. Impact of Mutations in the Limit Λ >> 1

In this section, we explain how the approach described in the main text needs to be amended in the presence of mutations. We focus here only on the case that Λ >> 1, because one can expect that the effect of mutations will be more dramatic in this limit corresponding to *hard* parasites. If one of such parasites is present in a compartment, it invades the population very quickly, provided it appears during the exponential growth phase. As explained in the main text, to describe the limit Λ >> 1, we can introduce three inoculation probabilities, p_ribo_ for compartments containing only ribozymes, *p_para_* for compartments containing parasites and *p_zero_* for empty compartments:

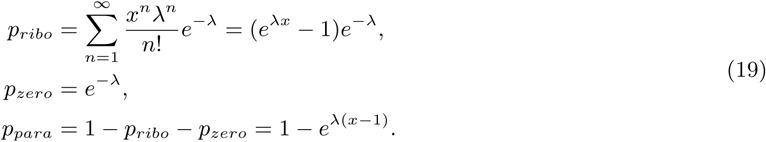

Now, let us also introduce *p_mut_* as the probability that a ribozyme is turned into a parasite as a result of a mutation during one replication event of the molecule. Then the probability that there is no mutation occurring during n* replication events is

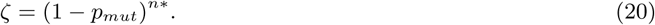

A typical value for this n* corresponds to what is denoted n_D_ in the previous section, namely the number of replications until the end of the exponential growth regime.

With *P_zero_* unchanged, the new probabilities for compartments containing ribozyme (resp. parasites) *P′_ribo_* (resp. P′_Para_) are simply

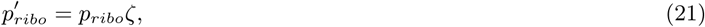

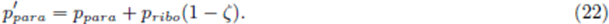

Using these expressions in the recurrence relation, we obtain

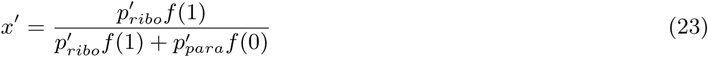

Evaluating the fixed point stability at *x* = 0 then yields the equation for the asymptote

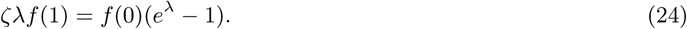

Similarly, we can evaluate the fixed point stability at *x* = 1, to obtain

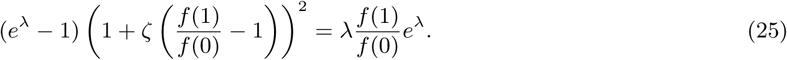

For *P_mut_ → 0, ζ → 1* and we obtain the asymptotes mentioned in the text. For ζ < 1, the asymptotic values of λ for both *x* = 0 and *x* = 1 become smaller. As a result, both the ribozyme and the bistable regions shrink as one would expect. In the extreme case where ζ → 0, both regions disappear completely since then the only solution to Eqs. (24-25) corresponds to λ = 0.

### IV. Asymptotic Behavior for λ → ∞

For large λ, for Λ close to 1 and *x* close to 1 (resp. 0), the most abundant compartments verify *m = n* or *m = n* − 1 (resp. *m* = 0 or *m* = 1). As λ is large, we can neglect fluctuations in *n* and we can take *n* = λ. We therefore only look at the recursion for a typical compartment with *n* = λ, with a simplified notation P*λ(n = λ, x, m*) = P*λ*(*x, m*), where

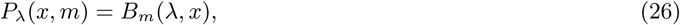

obtaining

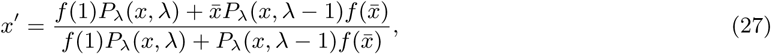

where

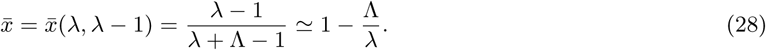

We have therefore

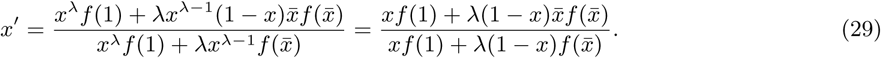

Taking the derivative with respect to *x* we obtain

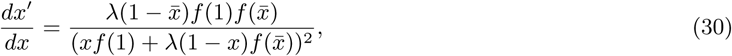

which for *x* = 1 yields

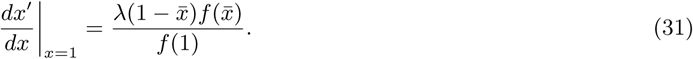

Thus the boundary defined by the equation

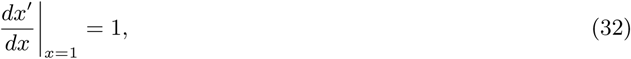

is given by

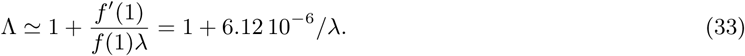

Evaluating the stability around the fixed point *x* = 0 we obtain likewise

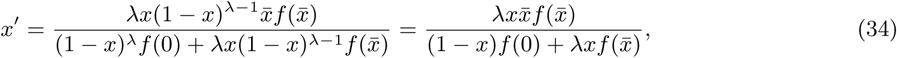

where now 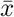 is given by

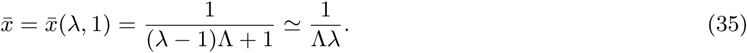

Evaluating the derivative of *x′(x)* at *x* = 0 we obtain

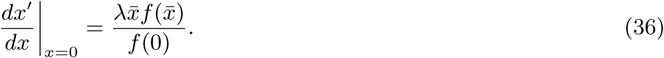

This gives the boundary as

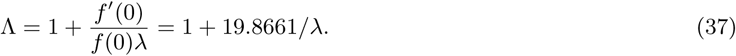

### V. Additional Features of the Phase Diagram

The construction of the phase diagram of the main text is based on the condition of stability of the two fixed points *x* = 0 and *x* = 1. This treatment only gives a complete picture of the phase behavior if there are at most three fixed points. While this is true for most pairs (λ,Λ), notice that in the special case of Fig. 8a for Λ = 4 and near λ = 8, the curve turns back. In this region, there are four fixed points, with *x* = 0 and 0 < *x*\* < 1 being stable. The novel aspect of the region near λ = 8 and *x* below 0.1 (shown in the inset as a blow-up), is that there is a bistability between points *x* = 0 and *x = x*\* whereas in the phase diagram of the main text, the bistability only concerned points *x* = 0 and *x* = 1. For *x* > 0.1 and λ between approximately 2 and 8, we have a standard coexistence phase.

**Fig. 8.**
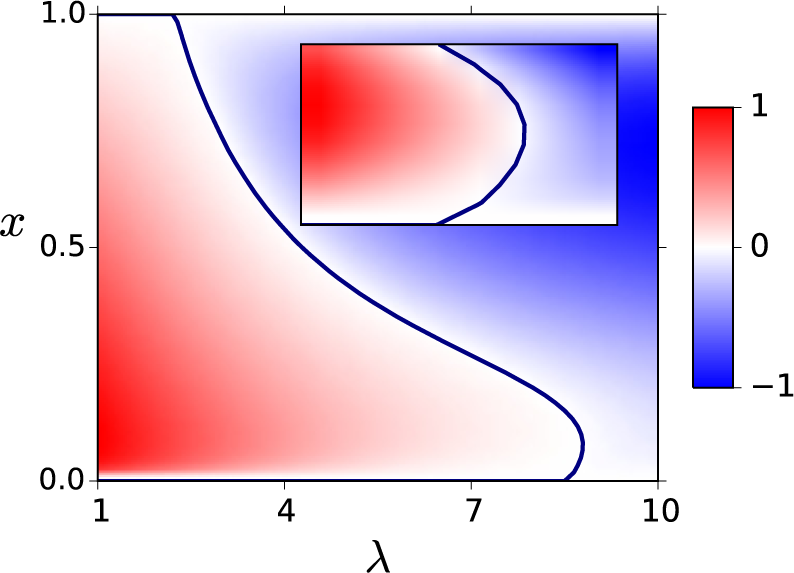
Contour plots of *Δx* vs. *x* for Λ = 4 in the plane (*x*, λ). Inset shows a blow-up of the region near λ = 8, which exhibits features of both the bistable and coexistence regions.

### VI. Comparison between Linear and non-Linear Selection Function

In the main text, we have introduced the following selection function

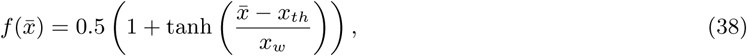

with *x*_th_ = 0.25 and *x_w_* = 0.1, which is now represented as the blue solid line in fig. 9. Note that this selection function takes a small but non-zero value for *x* = 0, namely 0.5(1 − tanh(*x_th_/x_w_*)) = 0.0067, which represents the fraction of false positives in the selection process.

**Fig. 9.**
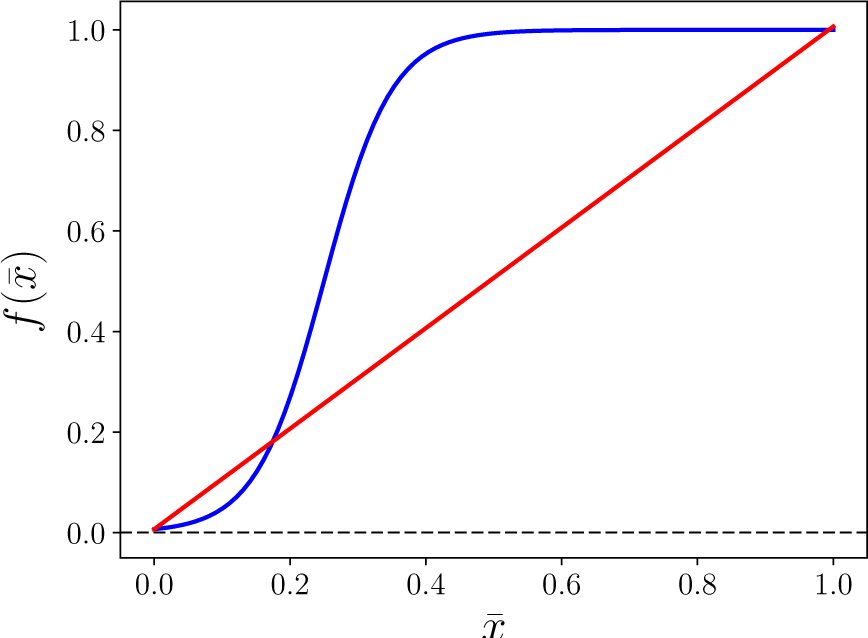
Representation of the two selection functions used in this section, namely the one denned in the main text (red solid line) and a linear one (blue solid line), which approximately take the same values at *x* = 0 and *x* = 1. The horizontal dashed line represents the level of false positives given by *f*(0).

**Fig. 10.**
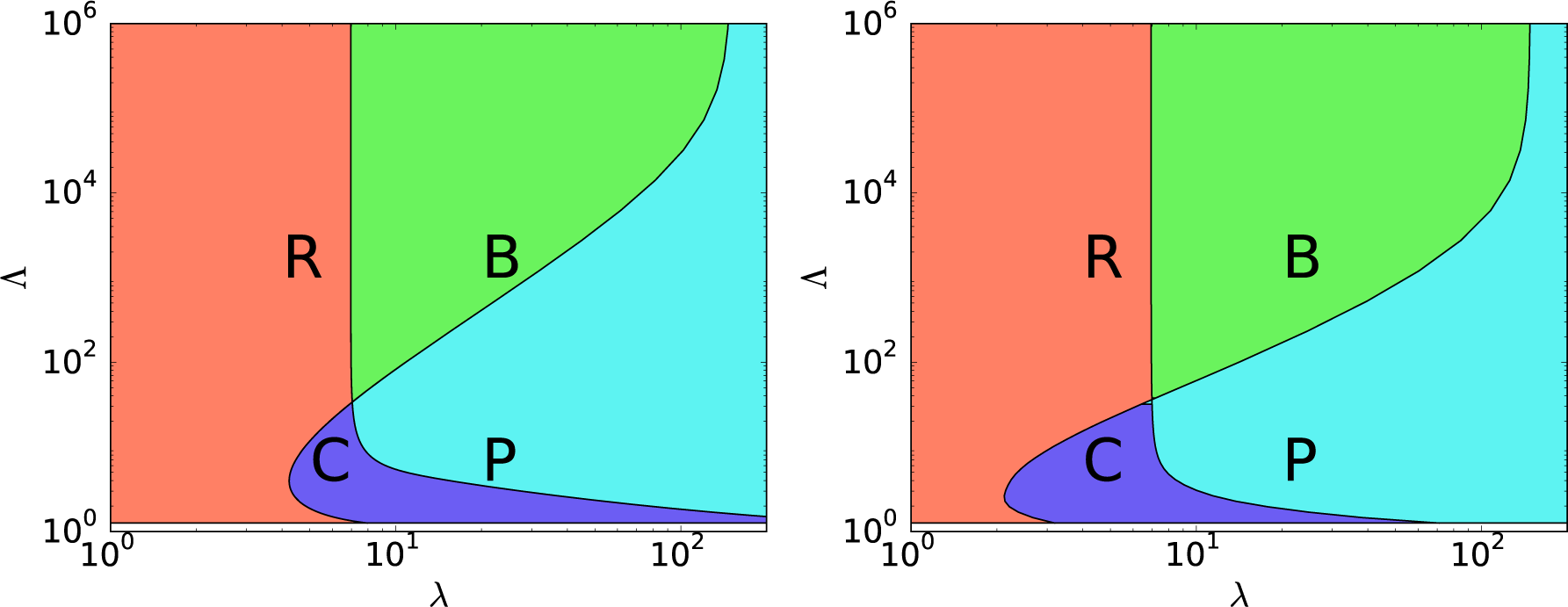
Left: Phase diagram of the transient compartmentalization dynamics with the linear selection function 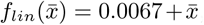in the (λ, Λ) plane. Right: idem with the non-linear selection function represented in Fig. 9. Phases are R: pure Ribozyme, B: Bistable, C: Coexistence and P: pure Parasite.

It is interesting to compare the phase diagram given in the main text with that obtained for a linear selection function, *f_lin_* shown as the red solid line in fig. 9. We choose 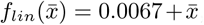, such that we have the same approximate values for f (0) and f (1) as with the previous function defined in Eq. (38). Consequently, we expect to find the same vertical asymptotes at λ ≃ 6.95 and λ ≃ 149, as confirmed in fig. 10. The equations of these vertical asymptotes are indeed only a function of f(0) and f(1). They are

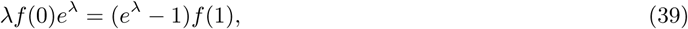

for the boundary between the bistable and parasite regions, and

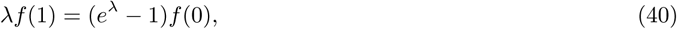

for the boundary between the bistable and ribozyme regions.

When comparing the two phase diagrams obtained with the non-linear and linear selection functions shown in fig. 10, we observe a similar general structure except for the center of the diagram and for the two asymptotes for Λ close to 1. This is to be expected for the center region where none of the simple approximations hold. Concerning the asymptotes near Λ = 1, as shown in the main text they represent boundaries between the coexistence and parasite regions (resp. coexistence and ribozyme regions) and they depend on the logarithmic derivative of the selection function near *x* = 0 (resp. *x* = 1).

